# Mitogen-activated protein kinases contribute to temperature induced cardiac remodelling in rainbow trout (*Oncorhynchus mykiss*) heart

**DOI:** 10.1101/2021.05.11.443523

**Authors:** Y. Ding, E.F. Johnston, T.E Gillis

## Abstract

Rainbow trout (*Oncorhynchus mykiss*) live in temperate environments and experience seasonal changes in temperature that range between 4°C and 20°C. Laboratory studies demonstrate that cold and warm acclimation of male trout can have oppositional effects on cardiac hypertrophy and the collagen content of the heart. The cellular mechanisms behind temperature induced cardiac remodelling are unclear, as is why this response differs between male and female fish. Recent work utilizing cultured trout cardiac fibroblasts suggests that collagen deposition is regulated, at least in part, by mitogen-activated protein kinase (MAPK) cell signalling pathways. We therefore hypothesized that temperature-dependent cardiac remodelling is regulated by these same cell signalling pathways. To test this, male and female trout were acclimated to 18°C (warm) in the summer and to 4°C (cold) in the winter and the activation of MAPK pathways in the hearts were characterized and compared to that of control fish maintained at 12°C. Animals, maintained under a natural photoperiod matched to time of year, were sampled throughout each acclimation. p38 MAPK phosphorylation increased in the hearts of female fish during the cold acclimation protocol and the phosphorylation of extracellular signal-regulated kinase (ERK) increased in the hearts of male fish with warm acclimation. These results indicate that thermal acclimation has transient and sex-specific effects on the phosphorylation of MAPKs.

## Introduction

An acute change in physiological temperature represents a challenge for any animal as it directly affects the function of all biochemical and biophysical processes. At the cellular level, for example, a reduction in temperature decreases enzyme activity as well as the fluidity of membranes. In the vertebrate heart, reduced temperature results in a loss in the rate and strength of contraction (Harrison and Bers, 1990). Despite the effects of temperature change on cardiac function, rainbow trout (*Oncorhynchus mykiss*) remain active in waters that seasonally range between 4°C and 20°C (Gamperl and Farrell, 2004; Keen and Farrell, 1994). It is hypothesized that these fish can compensate for the effects of temperature on cardiac function through the remodeling of tissue composition, morphology, and functional capacity (Gamperl and Farrell, 2004; Graham and Farrell, 1989; Keen et al., 2016; Klaiman et al., 2011; Klaiman et al., 2014, Aho and Vornanen, 2001). For example, cold acclimation has been found to increase the Ca^2+^ sensitivity of the trout myocardium, cause hypertrophy of the spongy myocardium, decrease the relative thickness of the compact myocardium, and induce cardiac fibrosis (Keen et al., 2016; Klaiman et al., 2011; Klaiman et al., 2014). Conversely, warm acclimation (17°C) of male trout can cause a decrease in relative ventricle size, an increase in relative compact layer thickness, and a decrease in extracellular collagen (Keen et al., 2016; Klaiman et al., 2011). It should be noted that not all cold acclimation studies report hypertrophy of the heart (Sephton and Driedzic., 1995; Driedzic et al., 1996; Gamperl and Farrell, 2004; Klaiman et al., 2014; and Keen et al., 2016). It has been proposed that it is not only temperature cues that initiate this response but also changes in photoperiod and cycling hormone concentrations including thyroid hormone and testosterone (Tiitu and Vornanen, 2003; Gamperl and Farrell, 2004, Klaiman et al., 2011). Such factors need to be considered when designing experiments and interpreting results.

The capacity to reduce collagen content in response to a physiological stressor is quite interesting considering that once collagen is deposited in the mammalian heart, it is permanent and plays a major role in progressive heart failure (Jalil et al., 1988, 1989; Marijianowski et al. 1995; Pauschinger et al. 1999). While the consequences of temperature induced cardiac remodeling have been described, the cellular pathways responsible for the alterations in collagen and muscle composition in fish are not yet known. In mammalian models, the deposition of collagen in the heart is mediated through the actions of angiotensin II (Ang II), and its induction of transforming growth factor-β (TGF-β) cytokine expression (Bernardo et al., 2010; Campbell and Katwa, 1997; Khan and Sheppard, 2006; Kim et al., 1995; Lee et al., 1995; Rosenkranz, 2004). Ang II and TGF-β bind to cell surface membranes and activate an intracellular signalling cascade that is received by the nucleus via a group of highly conserved signalling molecules called mitogen-activated protein kinases (MAPKs) (Kaschina and Unger, 2003; Moriguchi et al., 1999; Rosenkranz, 2004; Wenzel et al., 2001; Widmann et al., 1999; Yamaguchi et al., 1995). The activation of MAPKs, specifically the p38-ERK-JNK pathway, alters the transcription of genes related to extracellular matrix remodelling, including an upregulation of collagen 1α (*col1α*), downregulation of matrix metalloproteinases (*mmp-2, -9, -13*), and upregulation of tissue inhibitor of matrix metalloproteinases (*timp-2*) (Nagase et al., 2006; Silvipriya et al., 2015; Visse and Nagase, 2003). The end result is an increase in the collagen content of the ECM (Nagase et al., 2006; Silvipriya et al., 2015; Visse and Nagase, 2003). This fibrotic response can increase cardiac stiffness, and when in excess, can lead to diastolic dysfunction.

In rainbow trout, cold-induced cardiac hypertrophy and fibrosis can occur in parallel, and are accompanied by increases in the expression of gene transcripts for *timp2* and *col1α3*, and decreases in the expression of *mmp-2* and *mmp-13* (Keen et al., 2016). As in mammals, these transcriptional responses result in an increase in collagen deposition (Nagase et al., 2006; Visse and Nagase, 2003). Conversely, collagen removal in the trout heart with warm acclimation is accompanied by a decrease in *timp2* and *col1α3* expression and an increase in *mmp-2* expression (Keen et al., 2016). Importantly, treatment of cultured trout cardiac fibroblasts, the cells that produce collagen, with physiologically relevant concentrations of TGF-β1 causes an increase in collagen deposition, and similar changes in the expression of *mmp-2, -9, timp-2*, and *col1α1* gene transcripts, as occurs with cold acclimation (Johnston and Gillis, 2017). This indicates that the signaling pathways regulating ECM are conserved, at least in part, in vertebrates.

It has been previously hypothesized that increased biomechanical stimulation of the heart is responsible for inducing cardiac remodeling in cold-acclimated fish (Graham and Farrell, 1989; Keen et al., 2016), and may be linked to MAPK signaling. Increased levels of biomechanical stimulation, sensed *via* mechanically sensitive cell membrane receptors, are thought to result from the increase in cardiac stroke volume that occurs in trout with cold acclimation, as well as from the measured increase in plasma viscosity (Graham and Farrell, 1989; Keen et al., 2016). The increase in stroke volume compensates for the decrease in heart rate at low temperatures (Aho and Vornanen, 2001), while the increase in blood viscosity is caused by a decrease in the fluidity of the erythrocyte membranes (Singh and Stoltz, 2002). An increase in plasma viscosity increases shear stress within the heart and this has been shown to translate into cardiac hypertrophy and fibrosis in mammalian models (Devereux et al., 1987). In addition, an increase in blood viscosity increases vascular resistance, and this translates into higher levels of work performed by the heart, and therefore greater biomechanical stimulation of the myocardium (Farrell, 1984; Keen et al. 2016). Indeed, previous studies have demonstrated that exposure of the mammalian heart to increased biomechanical stretch triggers an increase in TGF-β1 and in collagen deposition (Katsumi et al., 2004; Mackenna et al., 1998). Furthermore, Johnston and Gillis (2020) found that exposure of cultured trout cardiac fibroblasts to cyclical, 10% equibiaxial stretch, to mimic the cardiac cycle, caused an increase in the activation of p38 MAPK and ERK 1/2, a response that would likely lead to an increase in collagen deposition. However, no one has examined if these cell signalling molecules are activated during cold acclimation *in vivo* by the impact of a change in temperature on the functional properties of the myocardium. If they are, they could be what links such effects to the cellular processes responsible for cardiac remodeling.

The purpose of the present study was to determine if thermal acclimation of rainbow trout activates the MAPK signalling pathway, which is hypothesized to be responsible for initiating temperature induced cardiac remodelling. Building from our *in vitro* studies (Johnston and Gillis, 2020; Johnston et al. 2019; Johnston and Gillis 2018; Johnston and Gillis, 2017), we specifically hypothesized that cold acclimation (4 °C) would cause an increase in the phosphorylation status of p38 MAPK and ERK, and induce changes in the expression of gene transcripts that would support an increase in collagen deposition. It was also hypothesized that warm acclimation (18 °C) would have the opposite effect. To test these hypotheses, rainbow trout were acclimated for 8 weeks to 4 °C in the winter or 18 °C in the summer and local photoperiod was maintained. For each experiment, control fish were maintained at 12 °C. Fish were periodically sampled to quantify p38 MAPK and ERK phosphorylation states in heart tissue along with transcript levels for *mmp -2, -9, -13* and *col1α3*. The fish utilized in the current study were kept in outdoor ponds prior to experimentation.

## Methods

### Animal husbandry

Mixed sex rainbow trout (O mykiss; n=288; ∼400-550 g) were obtained from Rainbow Springs Trout Farm (Thamesford, Ontario) where they were housed in outdoor ponds fed with groundwater at a constant 8°C throughout the year. Half of the fish were obtained in November for the cold acclimation experiment (December 2017-February 2018), and the other half in May for the warm acclimation experiment (June 2018-August 2018). For each acclimation experiment, fish were transported to the Hagan AquaLab (University of Guelph, Ontario), transferred into one of two 2000 L environmentally controlled aquatic recirculation systems (ECARS), and then maintained for 10 d at 12°C before initiating the thermal acclimation protocol (see below). One of these tanks was maintained at 12 °C for the entire experiment (control), while the temperature of the other was changed according to the acclimation protocol (experimental). This enabled each thermal acclimation experiment to have its’ own control. Throughout the study, the trout were fed approximately 2% of their body weight every second day and maintained under the local outdoor photoperiod (43°31’43“N 80°13’37”W). This means that the cold acclimation experimental fish experienced a winter photoperiod, while the warm acclimation experimental fish experienced a summer photoperiod. At the start of the cold acclimation, the average mass of the fish was 430 ± 22 g, and at the start of the warm acclimation the average mass of the fish was 543.6 ± 9 g. All animal utilization protocols were approved by the University of Guelph Animal Care Committee (AUP #3958).

### Thermal acclimation

For the cold acclimation, the temperature of the water in the experimental tank was decreased by 1°C per day until the temperature reached 4°C. This rate of temperature change has been used in multiple previous studies (Aho and Vornanen, 2001; Keen et al., 2016; Klaiman et al., 2011; Klaiman et al., 2014). For the warm acclimation, the temperature of the water in the experimental tank was increased by 1°C per day until the water temperature reached 18°C. For both acclimations, the temperature of the control tank remained at 12°C, and fish were held at the target temperature for 8 weeks. Each acclimation had 6 sampling timepoints. For the cold acclimation, trout were sampled once the experimental temperature reached 8°C (Week -1/2), 4°C (Week 0), and at 1, 2, 4, and 8 weeks after reaching 4°C (W1, W2, W4, W8 respectively). For the warm acclimation experiment, trout were sampled once experimental temperature reached 15°C (Week -1/2), 18°C (Week 0), and at 1, 2, 4, and 8 weeks after reaching 18 °C (W1, W2, W4, W8 respectively).

### Tissue collection

At each sampling point, 12 fish were randomly selected from the experimental and control tanks. The trout were euthanized by a rapid cranial blow and severance of the cerebral spinal cord. The length and weight of the fish were measured and then the intact heart was removed, washed by immursion in ice cold heparinized physiological buffered saline (in mM: 94 NaCl, 24 NaCO_3_, 5 KCl, 1 MgSO_4_, 1 Na_2_HPO_4_, and 0.7 CaCl_2_, 20 pH 7.6 at 15°C, 15 units/mL of heparin) and immersed in ice cold 1 M KCl to induce maximal contraction. The ventricle was dissected from each heart and weighed. A portion from the same middle area of each ventricle was set aside for histological measurement. The remainder of the ventricles were flash frozen with liquid nitrogen and then stored at -80°C until further analysis. The time from when fish was killed to when the heart was flash frozen was aproximalty 1.5 minutes. Fulton’s condition factor, K, a measure of leaness in salmonid fish, was determined for each fish sampled according to Barnham and Baxter, (1902) using the equation below. A smaller K value indicates that fish are more lean.

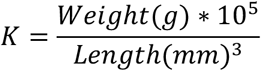

### Measurement of total and phosphorylated p38 and ERK1/2 with Western blotting

Antibodies specific to the MAPKs p38 and ERK1/2 and their phosphorylated forms were used to measure MAPK activation. Mouse cardiac tissue was used as a positive control for all of the antibodies used. All MAPK antibodies were obtained from Cell Signalling Technology (tot-ERK1/2: 4695S; p-ERK1/2: 4370S; tot-p38: 8690S; p-p38: 4511S; Danvers, Massachusetts). Western blotting procedures used were as described in Johnston and Gillis (2017). In brief, samples were diluted in 5x Laemmli buffer so that 20 μg of protein was loaded protein in each lane of the 12% polyacrylamide gels. Precision Plus Dual Colour (Bio-Rad) standard and an internal control were included on all gels. The internal control was created by pooling a random selection of samples. Once run, each gel was electro-blotted (Mini Trans-Blot Cell; Bio-Rad) onto a polyvinylidene fluoride membranes and then blocked in 5% bovine serum albumin in Tris-buffered saline with 0.1% Tween 20 (blocking buffer), followed by sequential incubations at 4°C in one of the primary antibodies (1:1000 in blocking buffer) and an anti-rabbit HRP-conjugated secondary antibody (1:2000 in blocking buffer (Cell Signaling Technology). Visualization of identified total protein and phosphorylated protein bands was completed using an Amershan ECL Plus detection kit (GE Healthcare) as per manufacturer’s instructions (Figure A1) and imaged using a ChemiDoc MP imaging system (Bio-Rad). Densitometry was conducted using ImageLab software (V. 6.0; Bio-Rad) and adjusted for the internal control. After visualization, membranes were stripped, and then stained with SYPRO Ruby (Molecular Probes) protein dye. Phosphorylated p38 and ERK1/2 were first standardized to total p38 and total ERK1/2. This was done to ensure that changes in phosphorylation that were detected were not a result of differential levels of the MAPKs within a sample. These standardized data were then normalized to total protein from the same membranes using a SYPRO Ruby Blot Stain (Bio-Rad). The averages of phosphorylated MAPK/total MAPK/total protein in the hearts from treatment fish, separated by sex, were compared to that from the hearts of time matched controls, separated by sex, using the statistical analyses outlined below.

### Measurement of type 1 collagen using Western blotting

Total type 1 collagen was measured in hearts collected at the beginning and end of each thermal acclimation (W0 and W8, respectively) and compared to control hearts. In brief, the collagen antibody (product #CL50171AP; Cedarlane, Burlington, Canada), was diluted to 1:1000 and proteins were separated on a 6% gel. When proteins were transferred from the gel to the membrane, 0.1% SDS was added to the transfer buffer. All other western blotting methods were identical as stated in the previous section.

### Measurement of gene expression using RT-qPCR

The transcript levels of the target genes, *col1a3, mmp-2, mmp-9, mmp-13*, were measured at multiple time points to establish a timeline of the cellular response during remodelling with respect to the signalling protein kinases and acclimation time. The primers used were those previously described by Johnston and Gillis (2018). *ef1α* and *β-actin* were determined to be the most suitable reference genes using the program GeNorm (Vandesompele et al., 2002). Hearts were processed as described in Alderman et al. (2017), then RNA was isolated and cDNA was synthesized as described in Alderman et al. (2018). RT-PCR was conducted as in Johnston and Gillis (2017). Transcript abundance was measured using a CFX96 Real-Time System (Bio-Rad) at default cycling conditions (annealing temperature set at 58.3°C) and a dissociation cycle. Each reaction contained 1x Power SYBR™ Green PCR Master Mix (ThermoFisher Scientific), 200 nmol of forward and reverse primer (Table 1), and 1:15 v:v of cDNA. The mRNA abundance of the genes was measured by fitting the threshold cycle value to standard curves prepared from serially diluted cDNA. All reactions generated a single-peaked melt curve. All transcript abundances were normalized to the geometric mean of the reference genes and standardized to the control group at each experimental time point. All non-reverse transcribed control samples failed to amplify.

### Histology

The middle portion of each ventricle was fixed overnight in 10% neutral buffered formalin and embedded in paraffin following standard procedures (Klaiman et al., 2014). Serial 5 μm transverse sections were collected on Superfrost slides (Fisherbrand) and heated in an oven for 1h at 100°C. Standard procedure for picrosirius red staining was followed to quantify collagen content (Rittié, 2017). In brief, sections were deparaffinized with xylene (2x, 3 min), and rehydrated with ethanol (100% 2x; 95%, 85%, 70%, 50% 1x; 3 min each). The sections were stained with picrosirius red stain then washed with 0.01 M HCl (1x, 2 min) and 70% ethanol (1x, 45 s). Finally, the sections were dehydrated with ethanol (95%, 100%; 1x, 3 min each) and cleared with xylene (2x, 3 min) before being mounted with Permount (Fisherbrand) and dried overnight.

### Imaging and quantification using ImageJ

Images of stained sections were taken with an inverted microscope at 40X magnification (Nikon Eclipse Ti; Nikon, Melville, NY, USA). 3-4 picrosirius-red stained sections per sample were imaged using brightfield and polarized light (Rittié, 2017). ImageJ (Fiji; Schindelin et al., 2012) was used to quantify collagen content as well as compact myocardium thickness. Collagen content was quantified on images taken with polarized light using the method described by Rich and Whittaker (2005). From this analysis, collagen hue was calculated and standardized to total collagen-staining pixels resulting in an average hue. This represents the degree of red staining collagen (i.e. thick/dense collagen) in the sample. Compact myocardium thickness was measured by taking thickness measurements (20 measurements per section) around the circumference of the section and standardizing to ventricle mass. All histological measurements were conducted on tissue dissected from the middle portion of the ventricle.

### Statistical methods

The effects of temperature and time on protein phosphorylation and gene transcript levels were considered separately for each sex using two-way ANOVAs followed by Bonferroni post-hoc tests where applicable. To determine the effects of seasonal photoperiod on collagen composition, proportional collagen fiber densities in all fish from the cold acclimation experiment (male and female, control and cold acclimated, sampled at Week 8; was compared to that of all fish from the warm acclimation experiment (male and female, control and warm acclimated, sampled at Week 8. This means that the compositional data for the control groups (male and female) and treatment groups (male and female) for the cold acclimation experiment were pooled together, and this was compared to the similarly pooled data from the warm acclimation experiment. We also tested for the effects of temperature acclimation on animal wet weight, Fulton’s condition index, relative atrium weight, relative ventricle weight, and relative compact layer thickness, at all sampling points using a Student’s t-test followed by a Bonferroni correction. Again we tested the means of each treatment, separated by sex, to that of the time matched controls, separated by sex. All data were tested for normality prior to analysis using a Shapiro-Wilk test. All statistical tests were performed with SigmaPlot (ver.12.5) and *p <* 0.05 was considered to be significant.

## Results

### p38 MAPK and ERK phosphorylation

In the cold acclimation experiment there was a significant main effect of time on the phosphorylation of p38 MAPK (F_5,64_=3.083, p=0.015) in the heart of male trout that was independent of treatment. Post-hoc tests revealed that p38 phosphorylation was significantly higher at W-1/2 than W0 and W8 (Fig 1A; p=0.044) (Fig 1A). In female trout, there was significant interaction between the effects of time and cold acclimation on p38 MAPK phosphorylation (F_5,55_= 7.122, p<0.001): whereas p38 MAPK phosphorylation was constant across timepoints in the control fish (p>0.05), cold-acclimated female trout had significantly higher p38 phosphorylation at W2 relative to other timepoints (p<0.001) and relative to control fish at W2 (p<0.001) (Fig 1B). There were no significant effects of time or temperature on ERK phosphorylation in either male or female trout (Fig 2A and 2B).

**Figure 1:**
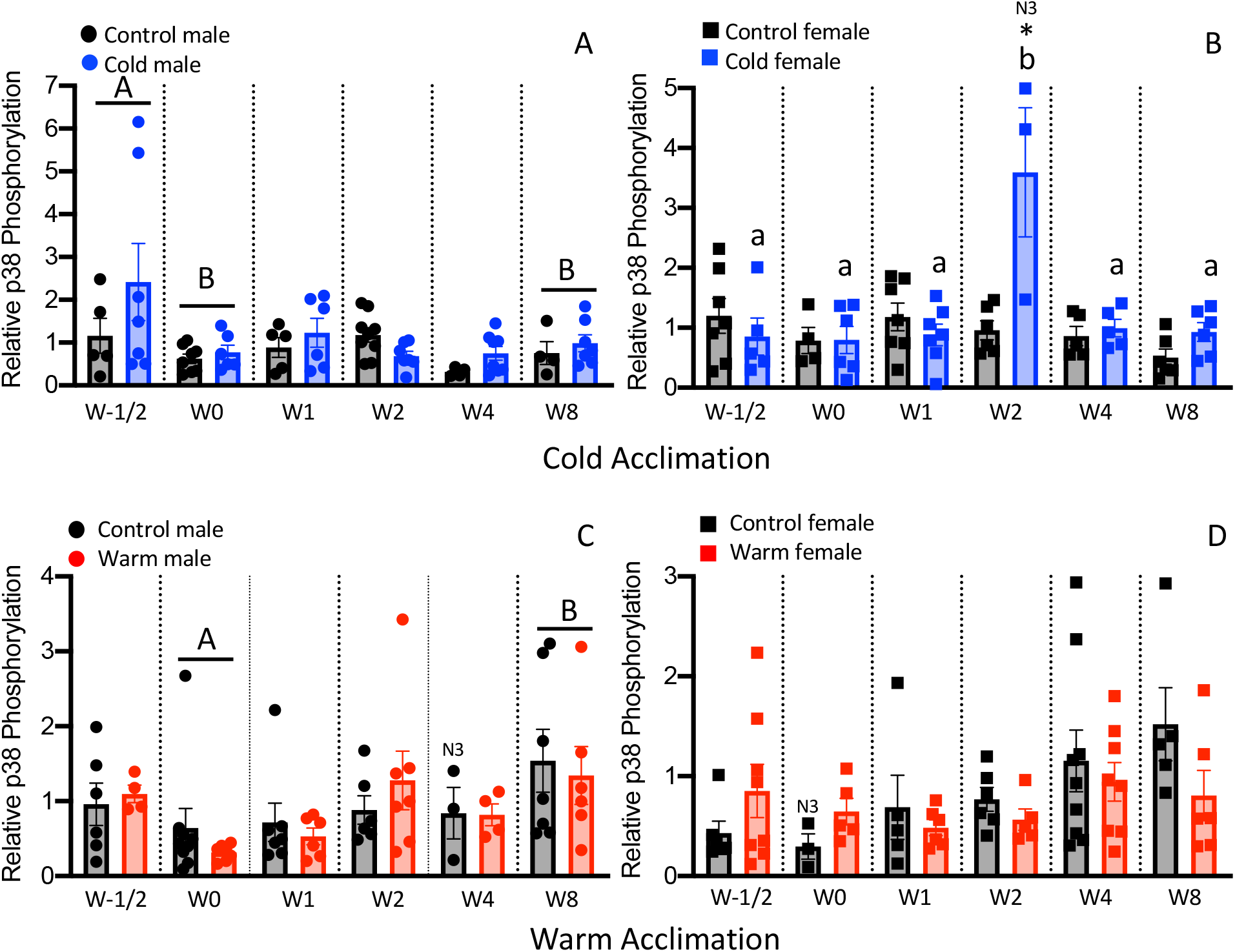
Relative p38 phosphorylation in the hearts of male and female rainbow trout (*Oncorhynchus mykiss*) across all sampling timepoints quantified with western blotting. Phosphorylated p38 was normalized first to total p38 then to total actin. A: Relative p38 phosphorylation in the heart of male trout during cold acclimation. B: Relative p38 phosphorylation in the heart of female trout during cold acclimation. C: Relative p38 phosphorylation in the heart of male trout during warm acclimation. D: Relative p38 phosphorylation in the heart of female trout during warm acclimation. W-1/2, halfway to final acclimation temperature 8°C (cold acclimation), 15°C (warm acclimation); W0, At final acclimation temperature 4°C (cold acclimation), 18°C (warm acclimation); W1, One week after acclimation; W2, Two weeks after acclimation; W4, Four weeks after acclimation; W8, Eight weeks after acclimation. N=3-9, N3 indicates a sample size of 3. * indicates significant difference between experimental and control fish (p<0.05). Bars annotated by different letters of the same case on a panel are different then each other (p<0.05).

**Figure 2:**
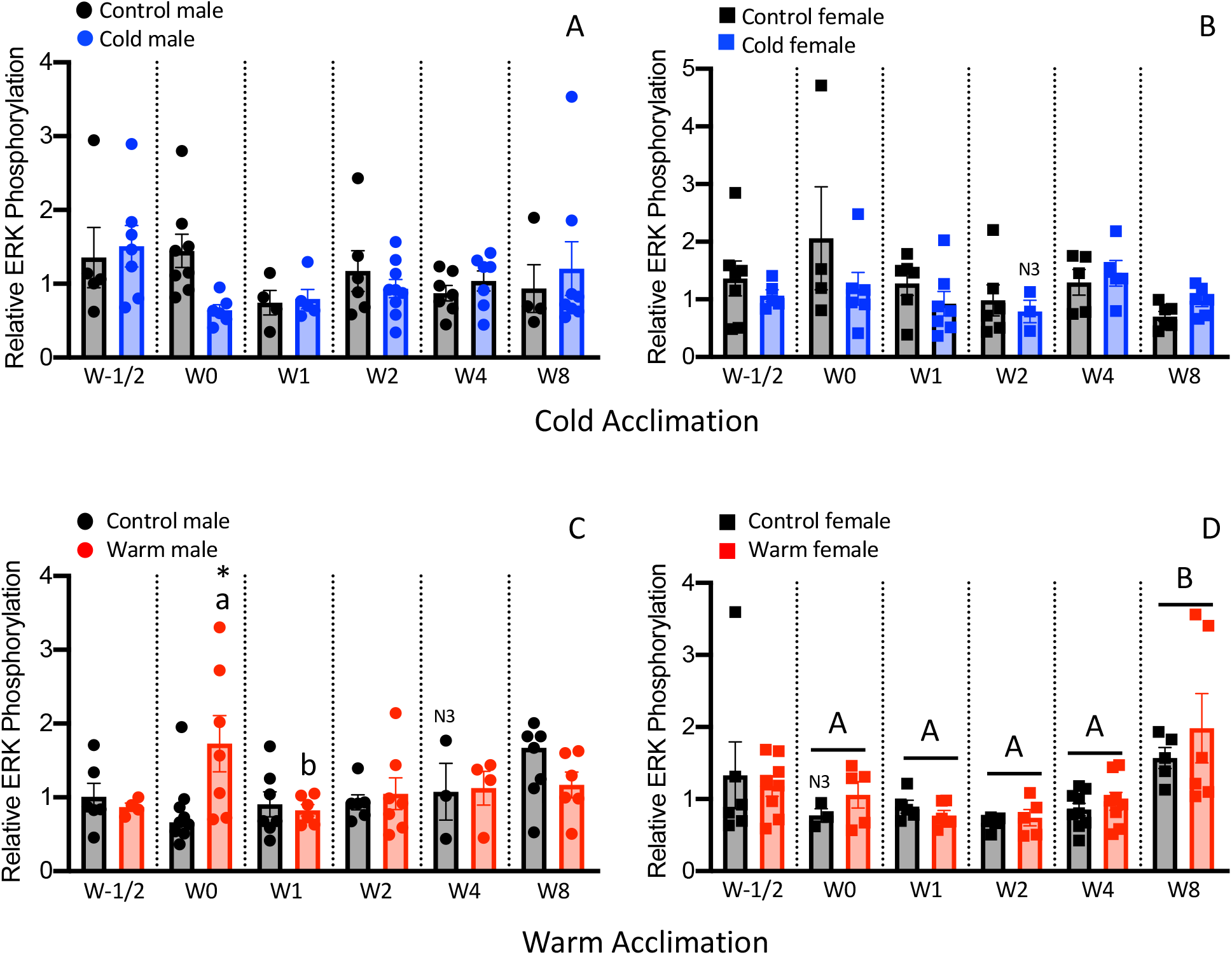
Relative ERK phosphorylation in the hearts of male and female rainbow trout (*Oncorhynchus mykiss*) across all sampling timepoints quantified with western blotting. Phosphorylated ERK was normalized first to total ERK then to total actin. A: Relative ERK phosphorylation in the heart of male trout during cold acclimation. B: Relative ERK phosphorylation in the heart of female trout during cold acclimation. C: Relative ERK phosphorylation in the heart of male trout during warm acclimation. D: Relative ERK phosphorylation in the heart of female trout during warm acclimation. W-1/2, halfway to final acclimation temperature 8°C (cold acclimation) 15°C (warm acclimation); W0, At final acclimation temperature 4°C (cold acclimation) 18°C (warm acclimation); W1, One week after acclimation; W2-Two weeks after acclimation; W4, Four weeks after acclimation; W8, Eight weeks after acclimation. N3 indicates a sample size of 3. * indicates significant difference between experimental and control fish (p<0.05). Bars annotated by different letters of the same case on a panel are different then each other (p<0.05).

In the warm acclimation experiment, there was a significant main effect of time on the phosphorylation of p38 MAPKs in the heart of warm acclimated male trout, (F_5,60_= 3.153, p=0.014). Post-hoc tests revealed that p38 phosphorylation was higher at W8 than W0 (Fig1C; p=0.011). There were no significant differences in p38 MAPK phosphorylation in warm acclimated female trout (Fig 1D). Two-way ANOVA revealed that there was significant interaction between time and treatment in warm acclimated male trout ERK phosphorylation (F_5,60_= 2.448, p=0.044). Post-hoc tests indicated that within W0, ERK phosphorylation was significantly higher in warm acclimated male trout than the control (Fig 2C; p<0.001) and that ERK phosphorylation was significantly higher in the warm acclimated trout at W0 than at W1 (Fig 2C; p<0.046) (Fig 1A). In warm acclimated female trout, there was a significant main effect of time on the phosphorylation of p38 MAPK (F_5,60_=2.452, p=0.044) and ERK (F_5,60_=5.820, p<0.001). Post-hoc tests did not find any significant differences in the levels of p38 phosphorylation between timepoints, but did reveal that ERK phosphorylation was significantly higher at W8 compared to W0, W1, W2, and W4 (p<0.02).

### *mmp -2, -9, -13* and *col1a3* transcript abundance

In the cold acclimation experiment, there was significant interaction between time and treatment in *mmp*2 transcript abundance in male cold acclimated trout (F_2,26_=9.871, p<0.001; Fig 3A). This was driven by a greater expression of *mmp2* at W8 in cold-acclimated male trout relative to earlier sampling times (p<0.002) and to W8 control (p<0.001). The transcript abundance of *mmp2* was also higher at W8 in female trout, but this was independent of cold acclimation (F_2,30_=4.819, p=0.015). In male warm acclimated trout, there was significant interaction between time and treatment on *mmp2* transcript abundance (F_2,30_=4.195, p=0.025), which was due to higher *mmp2* transcript abundance in control fish at W8 relative to warm acclimated male trout (p<0.001). There were no significant effects of time and treatment on *mmp2* transcript abundance in warm acclimated female trout.

**Figure 3:**
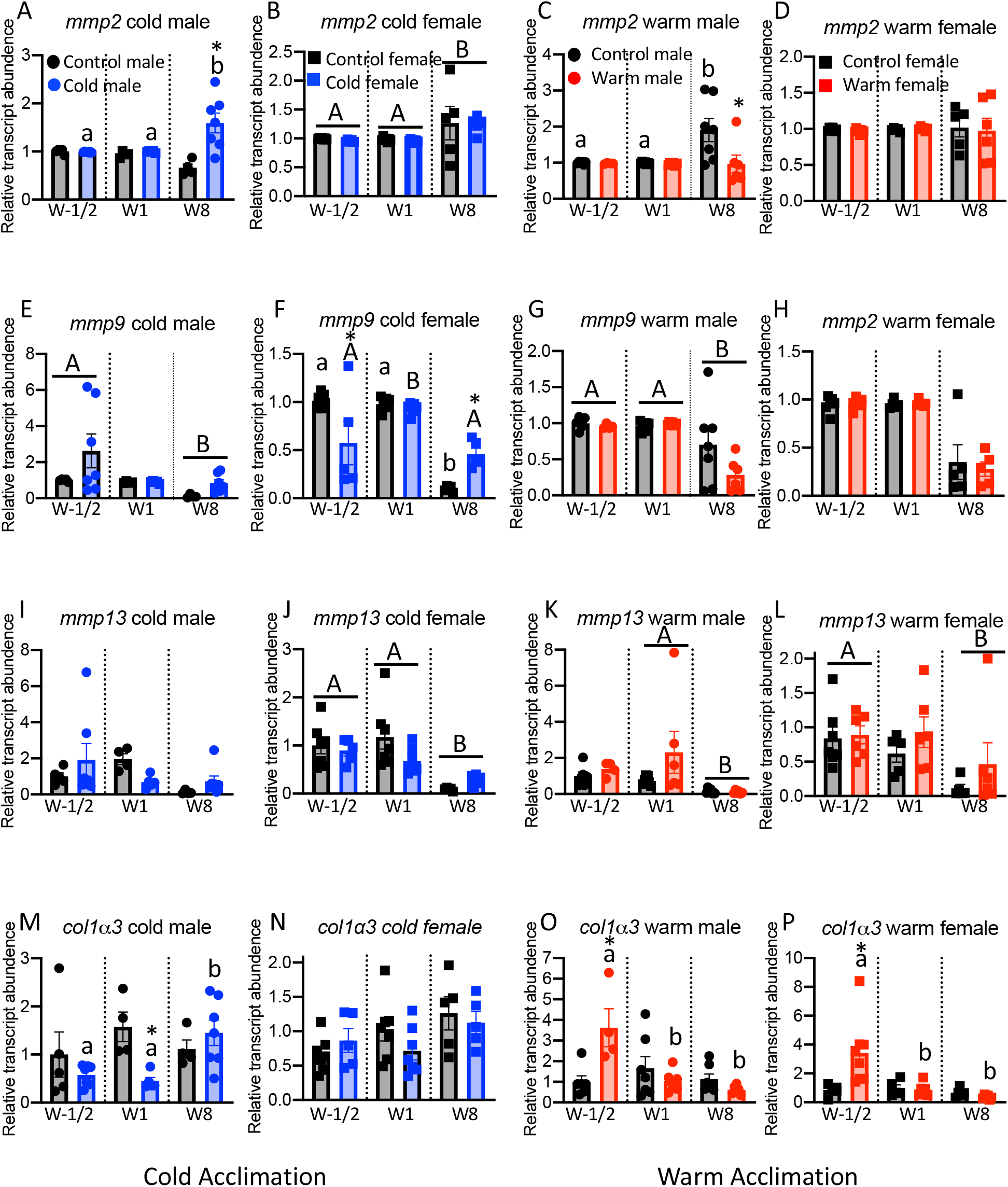
Influence of thermal acclimation on the relative transcript abundance of *mmp2, mmp9, mmp13*, and *col1α3* in the heart of male and female rainbow trout (Oncorhynchus mykiss). A) *mmp2* in cold acclimated male rainbow trout. B) *mmp2* in cold acclimated female rainbow trout. C) *mmp2* in warm acclimated male rainbow trout. D) *mmp2* in warm acclimated female rainbow trout. E) *mmp9* in cold acclimated male rainbow trout. F) *mmp9* in cold acclimated female rainbow trout. G) *mmp9* in warm acclimated male rainbow trout. H) *mmp9* in warm acclimated female rainbow trout. I) *mmp13* in cold acclimated male rainbow trout. J) *mmp13* in cold acclimated female rainbow trout. K) *mmp13* in warm acclimated male rainbow trout. L) *mmp13* in warm acclimated female rainbow trout. M) *col1α3* in cold acclimated male rainbow trout. N) *col1α3* in cold acclimated female rainbow trout. O) *col1α3* in warm acclimated male rainbow trout. P) *col1α3* in warm acclimated female rainbow trout. Transcript abundance was standardized to β-actin and ef1α. * indicates significant difference between experimental and control fish (p<0.05). N=4-7. Bars annotated by different letters of the same case on a panel are different then each other (p<0.05).

There was a significant effect of time on *mmp9* transcript abundance in male cold acclimated trout (F_2.26_=3.495, p=0.045). Post-hoc tests indicate that *mmp9* transcript abundance was lower at W8 than W1/2 (Fig 3C; p=0.049). In female cold acclimated trout, there was significant interaction between time and treatment on *mmp9* transcript abundance (F_2,30_=10.903, p<0.001) Within the female control group of the cold acclimation experiment, W8 was significantly lower than both W1/2 and W1. On the other hand, in cold acclimated female trout, *mmp9* transcript abundance was significantly higher at W1 than W1/2 and W8. Additionally, cold acclimation of female trout significantly decreased *mmp9* transcript abundance at W1/2 (p<0.001) and significantly increased *mmp9* transcript abundance at W8 (p=0.007). There was also a significant effect of time on the *mmp9* transcript abundance in male warm acclimated trout (F_2,30_=12.773, p<0.001). Post-hoc tests revealed that *mmp9* expression was significantly lower at W8 than W1/2 and W1 (Fig 3D; p<0.001). There was no significant effect of time or acclimation on the expression of *mmp9* in female warm acclimated trout.

The transcript abundance of *mmp13* was not affected by time or treatment in male cold acclimated trout. However, in female cold acclimated trout, time had a significant effect on *mmp13* transcript abundance (F_2,30_=12.878, p<0.001). Post-hoc tests revealed that the transcript abundance of *mmp13* was significantly lower at W8 compared to W1/2 and W1 (Fig 3E; p<0.001). Time also significantly impacted *mmp13* expression in the warm acclimation experiment in both males (F_2,30_=4.748, p=0.016) and females (F_2,29_=4.951, p=0.014). In males, post-hoc tests show that *mmp13* transcript abundance was higher at W1 than W8 (Fig 3F; p=0.017), while in females, transcript abundance was significantly higher at W1/2 than W8 (p=0.017).

There was significant interaction between the effects of time and treatment on *col1a3* transcript abundance in male cold acclimated trout (F_2,26_=3.602, p=0.042). Post-hoc tests indicate that at W1, *col1a3* expression was significantly lower in cold acclimated trout than the control (Fig 3G; p=0.009). Additionally, within the cold acclimated treatment group, *col1a3* expression was significantly greater at W8 than W1/2 and W1 (p<0.033). In female cold acclimated trout, time had a significant effect on the expression of *col1a3* (F_2,30_=3.364, p<0.048). However, post-hoc tests did not reveal any significant differences of transcript abundance between timepoints. In warm acclimated male trout, there was significant interaction between time and treatment on the expression of *col1a3* (F_2,30_=8.886, p<0.001 Post-hoc tests indicate that transcript abundance of *col1a3* was significantly greater at W1/2 than W8 in male warm acclimated trout (Fig 3H; p=0.006). Additionally, within W1/2, *col1a3* expression was significantly higher in the experimental group than the control (p<0.001). The *col1a3* transcript abundance of male trout exposed to the warm acclimation was significantly greater at W1/2 than W1 and W8 (p<0.001). In warm acclimated female trout, there was significant interaction between time and treatment (F_2,30_= 6.085, p=0.006), and this was due to greater transcript abundance of *col1α3* at W1/2 in warm acclimated females relative to later time points and to W1/2 controls.

### Collagen Western blots

There were no significant effects of time or acclimation on the collagen content of male or female rainbow trout (p>0.05). Figure A2.

### Collagen fiber thickness

Analysis of the polarized images of the microscopy sections from the two thermal experiments revealed that thick/dense (red) fibres were more prevalent in the hearts of fish used in the cold acclimation experiment (94.5 ± 4%) than in the hearts of fish used in the warm acclimation experiment (90.0 ± 5%) (*p<*0.05; Fig. 4). This was independent of treatment. Additionally, there were fewer orange fibres in the hearts of the fish used in the cold experiment than in the warm acclimation experiment (p*<*0.05). The only effect of thermal acclimation on collagen composition, detected using polarized light, was that there were fewer yellow (thin) fibres in the hearts of W8 female warm acclimated trout (4.4E-03±6.4E-04 %), than in the relevant control (1.6E-02±5.5E-03 %) (Table A1; *p*<0.05).

**Figure 4:**
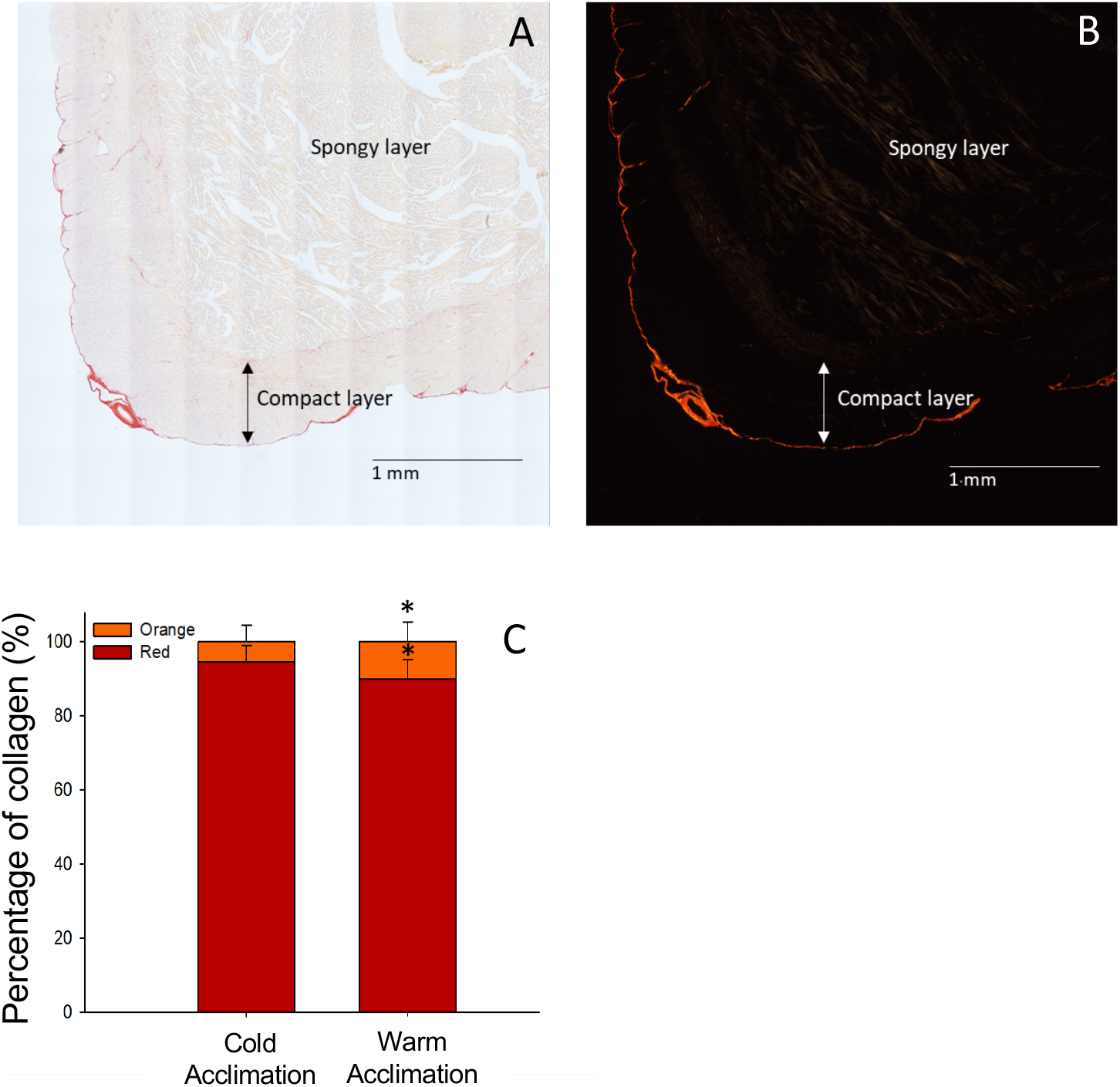
Histological analysis of ventricle collagen content. Representative bright field image (A) and polarized image (B) of ventricle section. (C) Relative proportion of collagen fiber olours in the ventricles of fish from the cold acclimation experiment vs. the warm acclimation experiment as quantified using picrosirius red imaging under polarized light. Red fibers represent the thickest fibers, while green fibers represent the thinnest fibres. Levels of green Cold 4.5e - 6%, Warm 3.6e-6%) and yellow (Cold 9.5e -5%; Warm 7.0e-5%) fibers were insignificant in comparison to the red and orange fibers and were therefore not plotted on the figure. The data for * indicates significant difference between the fish from the cold acclimation and warm acclimation experiments. N=4-7

### Growth and cardiac morphological measurements

There were no significant differences in body mass between the experimental fish and the relevant time matched controls at any of the sampling time points. Body condition also did not differ between experimental fish and controls at any time point. A summary of all morphological values can be found in Table A2. There was no difference in relative ventricle weight between thermally acclimated male and female fish and the relevant, control fish at any sampling timepoint. There was also no difference in the relative atrium weight between the male fish and relevant controls at any sampling point (Table A2). In the warm acclimation experiment, the relative atrium weight of the W8 female fish was significantly lower than that of control fish (Table A2; *p*<0.05). At all other timepoints, there were no significant differences. There was no difference in compact layer thickness relative to ventricle mass between any treatment fish (cold acclimation, warm acclimation) and the relevant time matched control (Table A2).

## Discussion

The results of this study deviate from what was predicted. While cold acclimation did result in an increase in relative p38 phosphorylation in the heart of female trout, warm acclimation also resulted in an increase in ERK phosphorylation in the heart of male trout. These changes in phosphorylation states suggest that thermal acclimation affects the p38-ERK-JNK cell signaling pathway; however the response is more complex than expected. The modest changes in the expression of gene transcripts associated with the regulation of collagen content indicate that thermal acclimation induced active regulation of the collagen content of the myocardium.

### Influence of thermal acclimation on p38 and ERK

The increased level of p38 phosphorylation with cold acclimation in the hearts of female trout at W2 suggests that a decrease in physiological temperature leads to an activation of this signalling molecule. We have proposed that the mechanism of action by which a decrease in physiological temperature leads to the activation of p38, as part of the MAPK signalling pathway, is *via* an increase in the stretch of the myocardium caused by an increase in stroke volume and blood viscosity (Johnston and Gillis, 2020). The results of this study support this proposal and build off the hypothesis by Graham and Farrell (1989) that an increase in blood viscosity, caused by a decrease in physiological temperature, is the trigger of cardiac remodeling in trout. Mechanically sensitive receptors located in the extracellular matrix (ECM), including integrins, have been shown to stimulate remodeling pathways within mammalian cardiac fibroblasts (Herum et al., 2017; Mackenna et al., 2000; Manso et al., 2009). Integrin subunits are associated with intracellular focal adhesion kinase, the phosphorylation of which is thought necessary for ERK and p38 signaling (Katsumi et al., 2004). In the heart, the transcriptional factors affected by the p38-JNK-ERK pathway are associated with cardiac remodeling (Pramod and Shivakumar, 2014; Sinfield et al., 2013). Although MAPK signaling is ubiquitous among all eukaryotic cells, fibroblasts are responsible for producing the components of the ECM, suggesting that MAPK signaling in cardiac fibroblasts would likely result in changes to extracellular protein production *via* altered gene expression (Johnston and Gillis, 2020). Related to this, we recently demonstrated that exposure of cultured trout cardiac fibroblasts to 20 min of physiologically relevant levels of cyclical stretch caused an increase in p38 phosphorylation, and that 24 hours of stretch resulted in the phosphorylation of p38 and ERK (Johnston and Gillis, 2020). This indicates that G-coupled protein membrane receptors in trout cardiac fibroblasts are mechanosensitive. Interestingly, in the current study, ERK phosphorylation levels in the hearts of male trout were found to increase with warm acclimation at W0. While the direction of change is opposite to what was predicted, this result suggests that thermal acclimation is a trigger for MAPK phosphorylation, but the response of the whole animal to the stimulus is more complex. One potential source of complexity is the presence of different cell types within the whole ventricle. Stretch treatment studies conducted on cultured rat myocytes observed increased phosphorylation of p38, ERK, and JNK (Seko et al., 1999). On the other hand, stretch treatment of rat cardiac fibroblasts increased ERK and JNK but not p38 MAPK phosphorylation (Mackenna et al., 1998). The MAPK signalling pathway is integral to a variety of intracellular functions that are not limited to cardiac remodelling (Cargnello and Roux, 2011; Chen et al., 2001; Widmann et al., 1999), and so it is possible that the observed MAPK activation occurred in cardiomyocytes and not fibroblasts. Thus, while MAPK signaling is known to regulate collagen deposition from fibroblasts *in vitro* (Nagase et al., 2006; Silvipriya et al., 2015; Visse and Nagase, 2003; Gillis and Johnston, 2017), the fact that downstream indicators of MAPK activation were not found in this *in vivo* study challenges the assignment of p38 phosphorylation to a specific cell type in the heart of cold-acclimated female trout.

### Influence of thermal acclimation on gene and protein expression

Review of the gene expression data collected over time indicates that there is inherent change in the measured transcript levels, that is independent of treatment. This is best illustrated by the pattern of *mmp2* expression in cold acclimated females where transcript levels in both treatment and control hearts are higher at W8 than at W-1/2 and W1. Such variation, suggestive of innate regulation of gene transcription, requires a conservative interpretation of results. The increased expression of *mmp2* in the heart of cold acclimated male trout at W8, if translated into an increase in MMP2 protein, would result in a greater capacity to degrade collagen. In addition, the lower level of *col1α3* at W0 would potentially translate into a decrease in collagen synthesis but the subsequent increase in the expression at W8 of this transcript would offset such a response. When considering how changes in the expression of gene transcripts involved in the regulation of collagen may affect its deposition, it is important to remember that it is the sum of the changes that determine collagen levels. For example, an increase in MMP protein expression could be offset, at least in part, by a concurrent increase in col1α3 expression. Likewise, any effect of the significantly low level of *mmp9* expression in the heart of female trout at W-1/2 on total collagen content would likely be reduced by the increase in the expression of this transcript at W1. However, the increase in expression of *col1α3* in the heart of both male and female trout at W-1/2 of the warm acclimation experiment suggests that these two groups are responding similarly to the treatment. As we did not sample prior to W-1/2, we do not know if there were changes in the phosphorylation of either ERK or p38 phosphorylation that could have initiated this change. Additionally, the lack of effect of either cold or warm acclimation on total collagen content when measured in hearts sampled at W8 suggests that the increase in col1α3 expression in both male and female hearts at W-1/2 had limited sustained consequence on cardiac remodeling.

The higher proportion of thick collagen fibers and corresponding lower proportion of thinner, less dense fibers in the hearts of fish on a winter photoperiod (cold acclimation experiment) sampled at W8 compared to that in the hearts of fish on a summer photoperiod (warm acclimation experiment) sampled at W8, could translate into the muscle having greater mechanical stiffness. Work by Keen et al. (2016) demonstrated that cold acclimation of trout caused an increase in the passive stiffness of the whole ventricle and this was linked to a measured increase in total collagen content. Previous studies have proposed that an increase in mechanical stiffness would provide greater structural integrity to the myocardium, and as a result, compensate for an increase in hemodynamic stress caused by a greater stroke volume and blood viscosity (Keen et al., 2017). As there was no difference in fiber composition between treatment and control for the same experiment in the current study, the difference between experiments is independent of acclimation temperature, but could reflect a seasonal response programmed by photoperiod. Timing of physiological changes to photoperiod is well established, and may help ensure that such changes are seasonally timed to a constant cue (Darrow et al., 1988; Kumar et al., 2010; Nakane and Yoshimura, 2019). Indeed, cold acclimation of trout has been found to cause less cardiac hypertrophy when the animals were kept on a 12:12 h light: dark photoperiod than on a natural photoperiod (Aho and Vornanen, 2001; Gamperl and Farrell, 2004; Keen and Farrell, 1994; Keen et al., 1993; Sephton and Driedzic, 1995).

### Cardiac morphology

Cold-induced changes to cardiac morphology are commonly reported in rainbow trout (Aho and Vornanen, 2001; Farrell et al., 1988; Graham and Farrell, 1989; Keen et al., 1993; Klaiman et al., 2011; Keen et al., 2016) and other fish species (Johnson et al. 2014), but variation in this response exists and warrants discussion. Even in our own work, we do not always observe clear cardiac hypertrophy in cold-acclimated rainbow trout across experiments (Klaiman et al., 2014). In the absence of changes to heart morphology with cold acclimation, trout may increase Ca^2+^ sensitivity of the cardiac myofilament as well as the rate and level of force generation by the ventricle (Klaiman et al, 2014). Such functional changes would help compensate for a decrease in cardiac contractility caused by a decrease in physiological temperature, while not being dependent on an increase in muscle mass.

There are a number of possible reasons for variability in the remodeling response between published studies, including the rather obvious influence of genetic background. Commercial aquaculture stocks are often acquired for laboratory studies on salmonids, but many such stocks have not been exposed to natural conditions for multiple generations and are the product of selective breeding programs. This is known to influence heritable traits including the stress response (Kittilsen et al., 2009). Interestingly, Johansen et al. (2011) have also demonstrated that familial differences in the stress response translates into differences in the cardiosomatic index (CSI). Secondly, there is also the potential effect of variation in environmental factors experienced by the fish prior to the experiment between studies influencing the response of the fish to the thermal acclimation. If the remodeling response is activated by seasonal environmental cues, a change in temperature is only one of these, another, as mentioned above, is photoperiod. It is for this reason that we purposely utilized fish in the current study that had been raised in outdoor ponds. In addition to variation among strains of the same species, an untested but interesting hypothesis is that varying levels of circulating hormones, including thyroid hormone, or testosterone, between experiments influence the biological response to thermal acclimation. In humans, for example, exercise-induced cardiac hypertrophy is more pronounced in men than in women and correlates with circulating testosterone levels (Ayaz and Howlett, 2015; Hanley et al., 1989; Merz et al., 1996). Similarly, we have previously found greater cardiac hypertrophy and remodeling in male trout with cold acclimation than in female trout. The levels of circulating sex hormones vary throughout the year in salmonids (Duston and Bromage, 1986; Pottinger, 1988; Goetz et al. 2011) and this would have the potential to influence the response of the heart to a physiological stress such as thermal acclimation.

### Sex specific responses

The results of the current study support sex specific responses to thermal acclimation as cold acclimation only caused an increase in p38 phosphorylation in the heart of female trout, while warm acclimation only caused an increase in ERK phosphorylation in the heart of male trout. Importantly, it has been established that differences in the strength and duration of MAPK activation greatly affects the impact of the signal (Kholodenko, 2006; Kholodenko and Birtwistle, 2009; Murphy and Blenis, 2006). Therefore, differences in the activation of the MAPK system have the potential to translate into significant differences in the remodelling response. This is perhaps reflected in the measured changes in gene expression where, except for the increase in expression of *col1δ3* in the heart of both male and female trout at W-1/2 of the warm acclimation experiment, there was little similarity in the gene expression pattern in the hearts of male and female fish with thermal acclimation.

### Perspectives

The results of multiple studies demonstrate that the trout heart is highly responsive to a change in physiological conditions, including temperature change, exercise training, and seawater acclimation, changing form and function to maintain its capacity to pump blood (Graham and Farrell, 1989; Dindia et al., 2017; Klaiman et al. 2011; Aho and Vornanen, 2001; Keen et al. 2016; Birjs et al. 2017). For animals living in an environment where conditions change predictably with the seasons, or that may begin to be affected by global climate change, this flexibility ensures the maintenance of heart function (Keen et al. 2016). The capacity to remodel the heart is dependent on the coordination of changes to multiple levels of biological organization (Keen et al. 2016). Work to understand this coordination is relevant not only to gain insight into the capacity of the heart to respond to a physiological stressor, but also of the mechanisms that drive pathological changes.

## SOURCES OF FUNDING

Y.D. was supported by an Ontario Graduate Scholarship. This work in the Gillis Lab is supported by a Discovery Grant, and a Discovery Accelerator Supplement, from the National Sciences and Engineering Research Council of Canada.

## APPENDIX

**Table A1:** Effect of cold and warm acclimation on the collagen composition of rainbow trout (Oncorhynchus mykiss) ventricles. A) Relative proportion of collagen hue in the ventricle. Colors form a gradient from thickest/densest fibers (Red) to the thinnest fibers (Green). B) The mean hue value of ventricles with a higher score indicating the presence of more thick fibers than thin fibers. Collectively, there was a significantly greater proportion of red collagen in the cold acclimation experiment (all groups combined) than the warm acclimation experiment. There was also a significantly lower proportion of orange collagen in the cold acclimation experiment than the warm acclimation experiment. * indicates significant difference between experimental and control groups (p<0.05).

| Hue | Cold Male |  | Cold Female |  | WarmMale |  | WarmFemale |  |
| --- | --- | --- | --- | --- | --- | --- | --- | --- |
|  | Ctrl | Expt | Ctrl | Expt | Ctrl | Expt | Ctrl | Expt |
| **Red (%)** | 92.2±4.2 | 95.6±0.96 | 94.5±1.3 | 94.7±1.7 | 90.0±1.7 | 89.6±2.4 | 88.0±3.0 | 91.9±2.0 |
| **Orange (%)** | 7.8±4.2 | 4.4±1.0 | 5.5±1.3 | 5.3±1.7 | 10.0±1.7 | 10.±2.4 | 12.0±3.0 | 8.1±2.0 |
| **Yellow (%)** | 9.3E-03±1.3E-03 | 5.7E-03±1.2E-03 | 9.7E-03±1.6E-03 | 1.5E-02±3.5E-03 | 4.3E-03±6.9E-04 | 4.9E-03±1.3E-03 | 1.6E-02±5.5E-03 | 4.4E-03±6.4E-04 * |
| **Green (%)** | 2.7E-04±1.1E-04 | 4.4E-04±1.7E-04 | 4.1E-04±1.5E-04 | 6.3E-04±8.2E-05 | 1.4E-04±4.6E-05 | 2.3E-04±1.0E-04 | 5.3E-04±3.8E-04 | 6.1E-04±4.8E-04 |
| **Collagen Hue** | 186.8±8.0 | 176.14±2.2 | 192.9±8.4 | 180.9±9.0 | 183.8±6.8 | 192.1±8.9 | 196.6±18.4 | 186.9±11.7 |

**Table A2:**
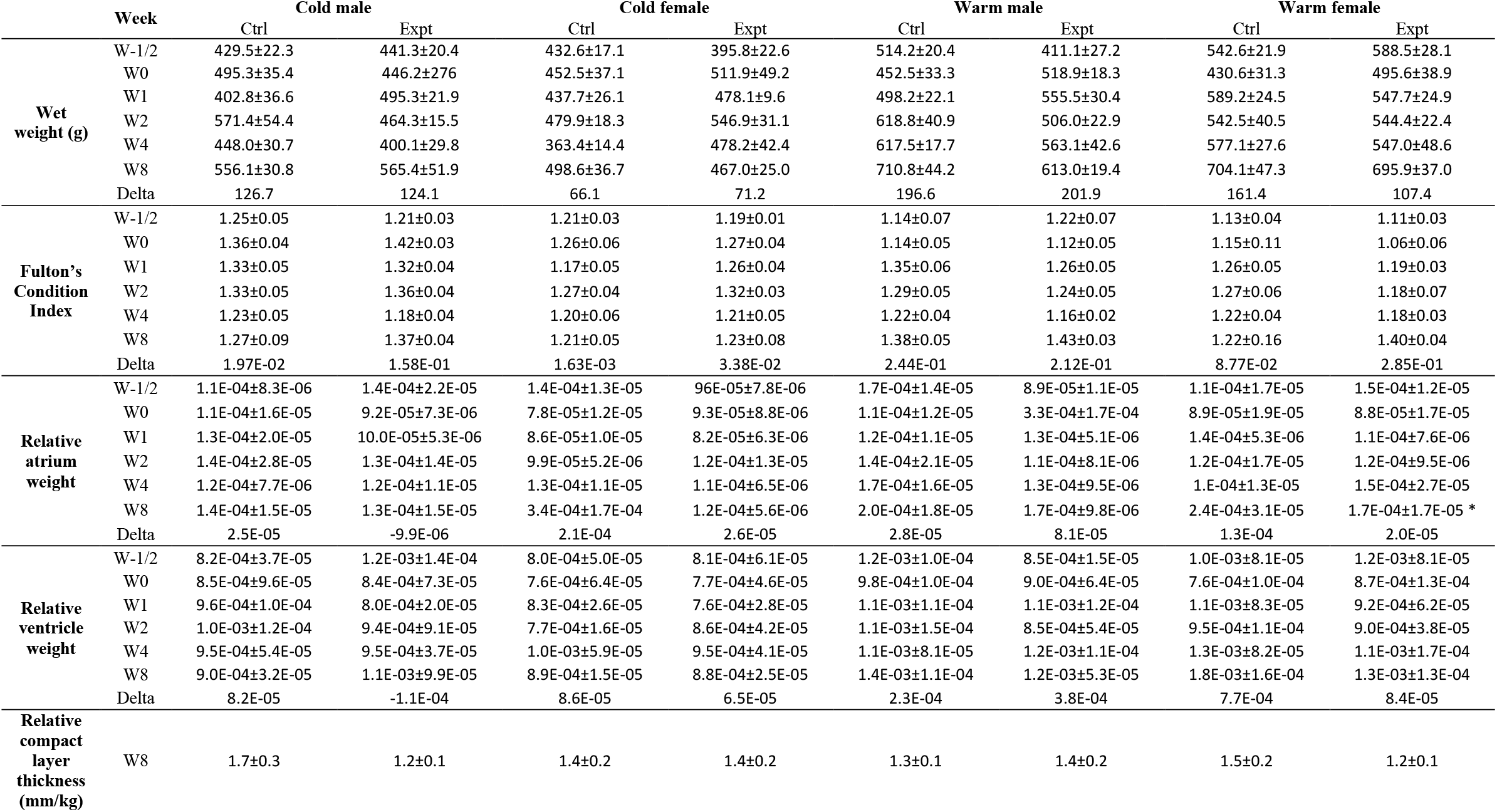
Summary of initial, final, and delta quantitative measurements (Mean ± SEM) taken during sampling. W-1/2-halfway to final acclimation temperature 8°C (cold acclimation) 15°C (warm acclimation); W0-At final acclimation temperature 4°C (cold acclimation) 18°C (warm acclimation); W1, One week after acclimation; W2, Two weeks after acclimation; W4, Four weeks after acclimation; W8, Eight weeks after acclimation. N=3-9. *indicates significant difference between experimental and control groups (*p*<0.05).

|  | Week | Cold male |  | Cold female |  | Warm male |  | Warm female |  |
| --- | --- | --- | --- | --- | --- | --- | --- | --- | --- |
|  |  | Ctrl | Expt | Ctrl | Expt | Ctrl | Expt | Ctrl | Expt |
| Wet weight (g) | W-1/2 | 429.5±22.3 | 441.3±20.4 | 432.6±17.1 | 395.8±22.6 | 514.2±20.4 | 411.1±27.2 | 542.6±21.9 | 588.5±28.1 |
| | W0 | 495.3±35.4 | 446.2±27.6 | 452.5±37.1 | 511.9±49.2 | 452.5±33.3 | 518.9±18.3 | 430.6±31.3 | 495.6±38.9 |
| | W1 | 402.8±36.6 | 495.3±21.9 | 437.7±26.1 | 478.1±9.6 | 498.2±22.1 | 555.5±30.4 | 589.2±24.5 | 547.7±24.9 |
| | W2 | 571.4±54.4 | 464.3±15.5 | 479.9±18.3 | 546.9±31.1 | 618.8±40.9 | 506.0±22.9 | 542.5±40.5 | 544.4±22.4 |
| | W4 | 448.0±30.7 | 400.1±29.8 | 363.4±14.4 | 478.2±42.4 | 617.5±17.7 | 563.1±42.6 | 577.1±27.6 | 547.0±48.6 |
| | W8 | 556.1±30.8 | 565.4±51.9 | 498.6±36.7 | 467.0±25.0 | 710.8±44.2 | 613.0±19.4 | 704.1±47.3 | 695.9±37.0 |
|  | Delta | 126.7 | 124.1 | 66.1 | 71.2 | 196.6 | 201.9 | 161.4 | 107.4 |
| Fulton's Condition Index | W-1/2 | 1.25±0.05 | 1.21±0.03 | 1.21±0.03 | 1.19±0.01 | 1.14±0.07 | 1.22±0.07 | 1.13±0.04 | 1.11±0.03 |
| | W0 | 1.36±0.04 | 1.42±0.03 | 1.26±0.06 | 1.27±0.04 | 1.14±0.05 | 1.12±0.05 | 1.15±0.11 | 1.06±0.06 |
| | W1 | 1.33±0.05 | 1.32±0.04 | 1.17±0.05 | 1.26±0.04 | 1.35±0.06 | 1.26±0.05 | 1.26±0.05 | 1.19±0.03 |
| | W2 | 1.33±0.05 | 1.36±0.04 | 1.27±0.04 | 1.32±0.03 | 1.29±0.05 | 1.24±0.05 | 1.27±0.06 | 1.18±0.07 |
| | W4 | 1.23±0.05 | 1.18±0.04 | 1.20±0.06 | 1.21±0.05 | 1.22±0.04 | 1.16±0.02 | 1.22±0.04 | 1.18±0.03 |
| | W8 | 1.27±0.09 | 1.37±0.04 | 1.21±0.05 | 1.23±0.08 | 1.38±0.05 | 1.43±0.03 | 1.22±0.16 | 1.40±0.04 |
|  | Delta | 1.97E-02 | 1.58E-01 | 1.63E-03 | 3.38E-02 | 2.44E-01 | 2.12E-01 | 8.77E-02 | 2.85E-01 |
| Relative atrium weight | W-1/2 | 1.1E-04±8.3E-06 | 1.4E-04±2.2E-05 | 1.4E-04±1.3E-05 | 96E-05±7.8E-06 | 1.7E-04±1.4E-05 | 8.9E-05±1.1E-05 | 1.1E-04±1.7E-05 | 1.5E-04±1.2E-05 |
| | W0 | 1.1E-04±1.6E-05 | 9.2E-05±7.3E-06 | 7.8E-05±1.2E-05 | 9.3E-05±8.8E-06 | 1.1E-04±1.2E-05 | 3.3E-04±1.7E-04 | 8.9E-05±1.9E-05 | 8.8E-05±1.7E-05 |
| | W1 | 1.3E-04±2.0E-05 | 10.0E-05±5.3E-06 | 8.6E-05±1.0E-05 | 8.2E-05±6.3E-06 | 1.2E-04±1.1E-05 | 1.3E-04±5.1E-06 | 1.4E-04±5.3E-06 | 1.1E-04±7.6E-06 |
| | W2 | 1.4E-04±2.8E-05 | 1.3E-04±1.4E-05 | 9.9E-05±5.2E-06 | 1.2E-04±1.3E-05 | 1.4E-04±2.1E-05 | 1.1E-04±8.1E-06 | 1.2E-04±1.7E-05 | 1.2E-04±9.5E-06 |
| | W4 | 1.2E-04±7.7E-06 | 1.2E-04±1.1E-05 | 1.3E-04±1.1E-05 | 1.1E-04±6.5E-06 | 1.7E-04±1.6E-05 | 1.3E-04±9.5E-06 | 1.E-04±1.3E-05 | 1.5E-04±2.7E-05 |
| | W8 | 1.4E-04±1.5E-05 | 1.3E-04±1.5E-05 | 3.4E-04±1.7E-04 | 1.2E-04±5.6E-06 | 2.0E-04±1.8E-05 | 1.7E-04±9.8E-06 | 2.4E-04±3.1E-05 | 1.7E-04±1.7E-05 * |
|  | Delta | 2.5E-05 | -9.9E-06 | 2.1E-04 | 2.6E-05 | 2.8E-05 | 8.1E-05 | 1.3E-04 | 2.0E-05 |
| Relative ventricle weight | W-1/2 | 8.2E-04±3.7E-05 | 1.2E-03±1.4E-04 | 8.0E-04±5.0E-05 | 8.1E-04±6.1E-05 | 1.2E-03±1.0E-04 | 8.5E-04±1.5E-05 | 1.0E-03±8.1E-05 | 1.2E-03±8.1E-05 |
| | W0 | 8.5E-04±9.6E-05 | 8.4E-04±7.3E-05 | 7.6E-04±6.4E-05 | 7.7E-04±4.6E-05 | 9.8E-04±1.0E-04 | 9.0E-04±6.4E-05 | 7.6E-04±1.0E-04 | 8.7E-04±1.3E-04 |
| | W1 | 9.6E-04±1.0E-04 | 8.0E-04±2.0E-05 | 8.3E-04±2.6E-05 | 7.6E-04±2.8E-05 | 1.1E-03±1.1E-04 | 1.1E-03±1.2E-04 | 1.1E-03±8.3E-05 | 9.2E-04±6.2E-05 |
| | W2 | 1.0E-03±1.2E-04 | 9.4E-04±9.1E-05 | 7.7E-04±1.6E-05 | 8.6E-04±4.2E-05 | 1.1E-03±1.5E-04 | 8.5E-04±5.4E-05 | 9.5E-04±1.1E-04 | 9.0E-04±3.8E-05 |
| | W4 | 9.5E-04±5.4E-05 | 9.5E-04±3.7E-05 | 1.0E-03±5.9E-05 | 9.5E-04±4.1E-05 | 1.1E-03±8.1E-05 | 1.2E-03±1.1E-04 | 1.3E-03±8.2E-05 | 1.1E-03±1.7E-04 |
| | W8 | 9.0E-04±3.2E-05 | 1.1E-03±9.9E-05 | 8.9E-04±1.5E-05 | 8.8E-04±2.5E-05 | 1.4E-03±1.1E-04 | 1.2E-03±5.3E-05 | 1.8E-03±1.6E-04 | 1.3E-03±1.3E-04 |
|  | Delta | 8.2E-05 | -1.1E-04 | 8.6E-05 | 6.5E-05 | 2.3E-04 | 3.8E-04 | 7.7E-04 | 8.4E-05 |
| Relative compact layer thickness (mm/kg) | W8 | 1.7±0.3 | 1.2±0.1 | 1.4±0.2 | 1.4±0.2 | 1.3±0.1 | 1.4±0.2 | 1.5±0.2 | 1.2±0.1 |

**Figure A1.**
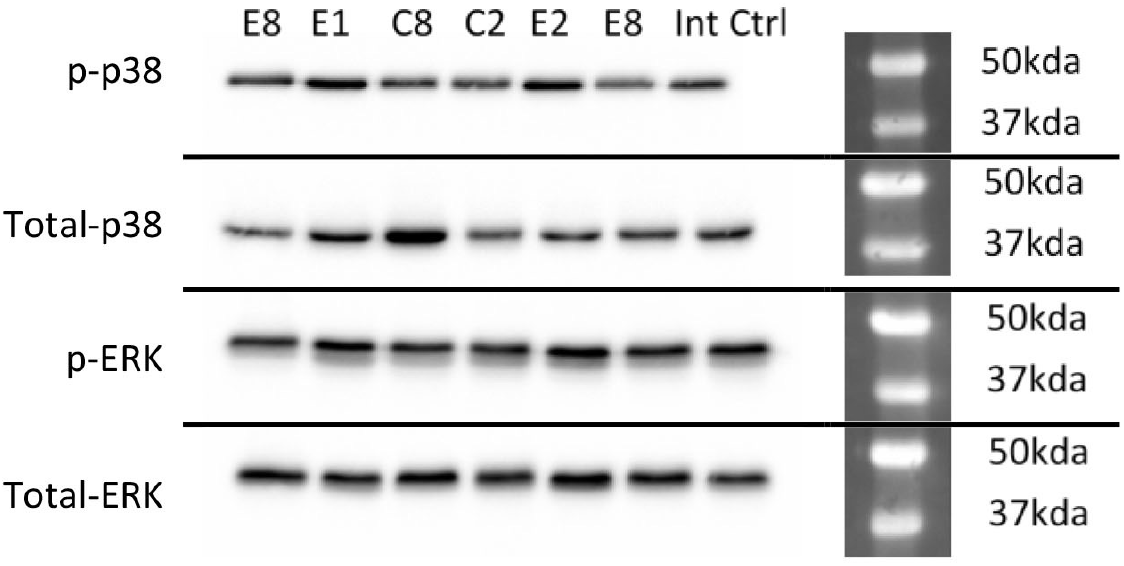
Representative western blot from the cold acclimation experiment. Each different antibody corresponds to different blots that were loaded and run simultaneously. Antibodies were from Cell Signalling Technology and are as follows: phosphorylated p38 (p-p38), total p38, phosphorylated ERK (p-ERK), total ERK. The sample group of each lane is listed along the top: E, Experimental; C, Control; Numbers indicate the sampling week (E8, experimental group sampled at W8). The same internal control sample was loaded with every blot.

**Figure A2:**
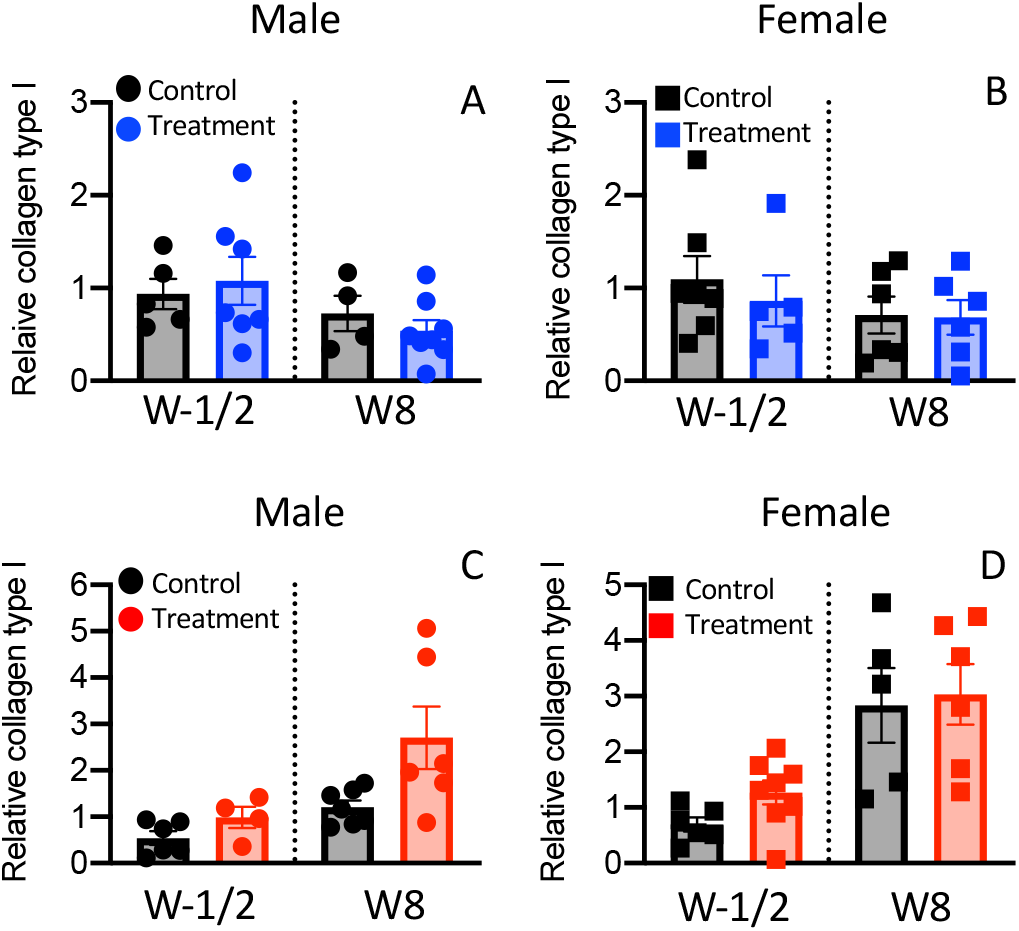
Relative ventricular collagen content, standardized to actin, measured by western blotting in rainbow trout (*Oncorhynchus mykiss*) at the initial (W-1/2) and final (W8) sampling periods. (A) Cold acclimated male heart, (B) Cold acclimated female heart. (C) Warm acclimated male heart, (D) Warm acclimated female heart. No statistically significant differences found (N=4-8). There are no statistically significant differences between the plotted means.

## Notes

### Competing Interest Statement

The authors have declared no competing interest.

